# The impact of commercially available ale and lager yeast strains on the fermentative diversity of beers

**DOI:** 10.1101/2020.07.17.209171

**Authors:** Diego Bonatto

## Abstract

Yeasts from the species *Saccharomyces cerevisiae* (ale yeast) and *Saccharomyces pastorianus* (lager yeast) are the main component of beer fermentation. It is known that different beer categories depend on the use of specific ale or lager strains, where the yeast imprint its distinctive fermentative profile to the beer. Despite this, there are no studies reporting how diverse, rich, and homogeneous the beer categories are in terms of commercially available brewing yeast strains. In this work, the diversity, richness, and evenness of different beer categories and commercial yeast strains available for brewing were evaluated by applying quantitative concepts of ecology analysis in a sample of 121,528 beer recipes. For this purpose, the frequency of ale or lager and dry or liquid yeast formulations usage was accessed and its influence in the fermentation temperature, attenuation profile, and number of recipes for a beer category were analyzed. The results indicated that many beer categories are preferentially fermented with dry yeast strains formulations instead of liquid yeasts, despite considering the high number of available liquid yeast formulations. Moreover, ale dry strains are preferentially used for lager brewing. The preferential use of specific yeast formulations drives the diversity, richness, and evenness of a beer category, showing that many yeast strains are potentially and industrially underexplored.

## 1. Introduction

Beer, a major alcoholic beverage obtained from malt-derived worts, is the product of fermentative metabolism of yeast strains that convert the sugars present in the wort into ethanol and CO_2_ (Rai and Jeyaram, 2017). The flavor impact of a specific yeast strain during beer fermentation is also important; in fact, many of the flavors found in a glass of beer are derived from metabolic by-products released by yeast cells during fermentation, like esters, lactones, thiol compounds, higher alcohols, and phenolics (Carrau et al., 2015; Praet et al., 2012; Tran et al., 2015).

Additionally, yeasts convert hop and malt-derived glycosylated metabolites to aglycones by the action of β-D-glucosidases during beer fermentation (Gamero et al., 2011); also, yeasts biotransform small molecules found in wort (e.g., amino acids and fatty acids) into flavor components (Carrau et al., 2015). Besides flavor, the visual aspects of a beer category are directly influenced by the yeast strains used for fermentation. For example, the clarity of the beer is a consequence of the flocculation ability of a yeast strain (Vidgren & Londesborough, 2011), while beer foam stability is also dependent on a series of glycoproteins present in the surface of yeast cell wall (Blasco & Viñas, 2011). Thus, the quality of beer is directly dependent on the yeast strain used.

Since the isolation and development of brewing yeast pure cultures from the works of Emil Christian Hansen in the end of 19^th^ century (Lodolo et al., 2008; Rank et al., 1988), and the identification of the yeast species that are responsible for bottom (lager) beer fermentation and top (ale) beer fermentation, the brewing industry has benefited from the use of yeast monocultures to give reproducible and consistent products over time. Two major yeast monocultures are employed in breweries nowadays, which are the *Saccharomyces cerevisiae*, mainly responsible for ale fermentation, and *Saccharomyces pastorianus*, a hybrid species responsible for lager fermentation (Lodolo et al., 2008). In this sense, cellular and molecular techniques are allowing researchers to design lager yeast strains for breweries (Mertens et al., 2015) and there is potential for the use of conventional (*S. cerevisiae*) and non-conventional yeast strains (e.g., *Saccharomyces eubayanus*) isolated from different environments niches for the design of new beers (Cubillos et al., 2019; Marongiu et al., 2015). Thus, the development of new hybrid strains or the use of environmental isolated yeast strains allow the brewer to explore different metabolic pathways and aggregate flavor diversity to beer (Cubillos et al., 2019). However, the applicability of new yeast strains in brewing industry could be impaired due to the genome and phenotype instabilities induced by the high selective and specific conditions of beer fermentations (Gorter de Vries et al., 2019), and brewers preferentially employ commercial yeast strains for beer fermentation due to the high fermentation efficiency and control (Bellissimi & Ingledew, 2005).

Therefore, considering the commercial available yeast strains for brewing it can be asked how diverse, rich, and homogeneous beer categories are in terms of different ale and lager yeast usage found in both dry and liquid formulations. For this purpose, a quantitative ecology analysis of diversity, richness, and evenness of commercial brewing yeast usage in different beer categories was performed by considering a sample of 121,528 beer recipes obtained from Brewer’s Friend web site (https://www.brewersfriend.com). In addition, the influence of fermentative parameters (e.g., lower and higher recommended fermentation temperature, and attenuation), yeast type (ale or lager), and formulation (dry or liquid) of commercial yeast strains used in beer categories fermentation were evaluated. The data gathered showed that beer categories can be classified as “cold fermented” and “hot fermented” considering the fermentation temperature profile of commercial yeast strains. Additionally, it was observed that there is a preferential use of dry yeast strains formulations for beer fermentation instead of liquid strains, even considering the high number of commercial yeast strains available in liquid formulations. Finally, it was observed that the preferential use of specific yeast type and/or formulation impacts the diversity, richness, and evenness of a beer category fermentative profile.

## 2. Material and methods

### 2.1. Commercial yeast strain data prospection and analysis

Data regarding yeast strains commercially available for breweries were obtained from Brewer’s Friend (https://www.brewersfriend.com; last access on May, 2020) with the direct consent of the web page administrator. Initially, the Lynx web browser (https://lynx.browser.org) was used to map all links associated with commercial yeast data strains, recipes, and different beer categories from Brewer’s Friend. Once obtained, the library rvest (https://github.com/tidyverse/rvest) from R software (https://www.r-project.org) was used to scrap recipe and commercial yeast data information for different beer categories from Brewer’s Friend links. The raw yeast and recipe data obtained were filtered and commercial yeast formulations containing the keywords “Wilds & Sours”, “Wine”, “S. boulardii”, “Mead”, “Cider”, “Champagne”, “Bretts and Blends”, “Bacterial Cultures”, “B. bruxellensis”, “Sake”, “Sour”, “Brett”, “Bug”, “Lactobacillus”, “Blend”, and “Saccharomycodes ludwigii” were removed from data. The resulting filtered yeast data containing information about manufacture company/laboratory, yeast strain brand name, type (ale or lager), formulation (dry or liquid), alcohol tolerance, flocculation, attenuation percentage, and lower and higher fermentation temperatures (in °C) were merged with beer category information. Finally, the definitions of beer categories as well as the country or geographical region from which they originated were obtained from the 2015 Beer Judge Certification Program (BJCP) Style Guidelines (https://dev.bjcp.org).

### 2.2. Statistical and quantitative ecology data analysis and preferential use of yeast strains

The R software (https://www.r-project.org) was used for all statistical and quantitative ecology data analysis. Data normality for quantitative variables for each beer category was evaluated by univariate Shapiro-Wilk normality test implemented in rstatix library (https://cran.r-project.org/web/packages/rstatix/index.html). Correlations between the number of yeast strains, recipes, lower and higher values of original and final gravity (OG and FG, respectively), international bitter units (IBUs), and alcohol by volume (ABV) were analyzed with corrplot library (https://github.com/taiyun/corrplot) by applying Spearman’s ρ statistic. All correlations with a *p-*value < 0.05 were considered statistically significant and were classified as follow: |*r*| = 0, null; 0 < | *r*| ≤ 0.3, weak; 0.3 < |*r*| ≤ 0.6, regular; 0.6 < |*r*| ≤ 0.9, strong; 0.9 < |*r*| < 1.0, very strong; |*r*| = 1.0, perfect. The library ggstatsplot (https://cran.r-project.org/web/packages/ggstatsplot/index.html) was used for comparing and plotting the lower and higher fermentation temperature as well as the attenuation percentage for brewing ale and lager yeasts strains in both dry and liquid formulations with the following parameters: display significant pairwise comparisons, Yuen’s method for robust estimation and hypothesis testing (Yuen, 1974), display confidence interval (CI_95%_) and estimated average value (μ), pairwise display all, evaluation of pairwise significance comparison by exact p-value, and false discovery rate adjustment method for *p-*values.

In order to evaluate the impact of the number of recipes in the lower and higher fermentation temperatures (LF_T_ and HF_T_, respectively) of a beer category, a weighted arithmetic mean value was determined considering the number of recipes for each beer category and the total number of recipes gathered from Brewer’s Friend web page using the following equations (1 and 2):

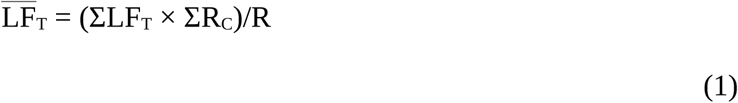

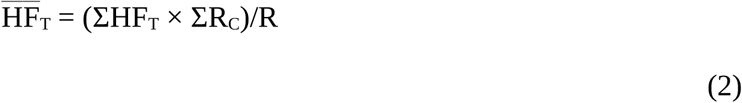

where 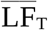 and 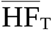 are the weighted arithmetic mean values for the lower and higher fermentation temperatures for each beer category, ΣLF_T_ and ΣHF_T_ represent the sum of lower and higher fermentation temperatures, respectively, for a given beer category, ΣR_C_ is the sum of the number of beer recipes for a given beer category, and R represents the total number of recipes available in Brewer’s Friend web page as obtained in May, 2020. Beer categories that display 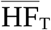 values above the average were classified as “hot fermented” beers, while beer categories with 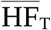 values below the average were classified as “cold fermented” beers. A linear regression analysis was performed in order to determine the correlation of 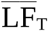 and 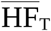 in different beer categories with the library ggpmisc (https://cran.r-project.org/web/packages/ggpmisc/index.html).

The preferential use of a specific brewing yeast strain (ale dry or liquid, and lager dry or liquid) in comparison to all different yeast strains reported for a beer category (*P*_*Y*_) was calculated as follow (equation 3):

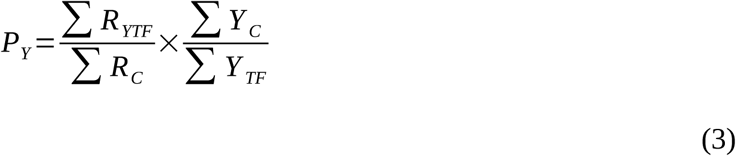

where Σ*R*_*YTF*_ is the total number of beer category-associated recipes that use a specific brewing yeast strain (ale dry or liquid, and lager dry or liquid), Σ*R*_*c*_ and Σ*Y*_*c*_ are the total number of recipes and yeast strains for a given beer category, respectively, and Σ*Y*_*TF*_ is the total number of beer category-associated specific brewing yeast strain (ale dry or liquid, and lager dry or liquid).

Quantitative ecology data analysis was performed in R environment with the vegan library (Dixon, 2003). In this sense, the frequency of a unique yeast strain in a beer category was used to estimate the parameters of richness, diversity, and evenness. For richness estimation, the Menhinick index (*Mi*) (Cazzolla Gatti et al., 2020) was applied with the equation (4):

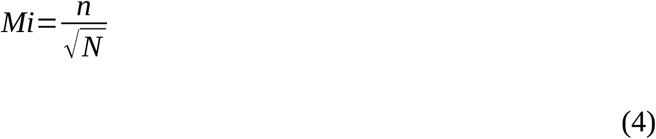

where *n* is the frequency of unique yeast strains for a given beer category and *N* is the number of recipes for a beer category. By its turn, the Simpson’s diversity (*D*^*S*^) (Thukral, 2017) of brewing yeast strains in different beer categories was determined by using the Simpson’s index (λ) described in equations 5 and 6:

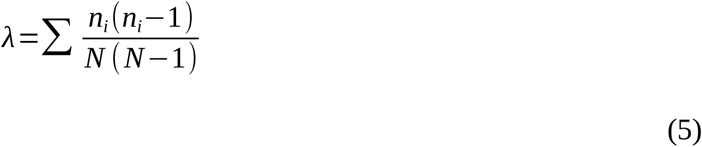

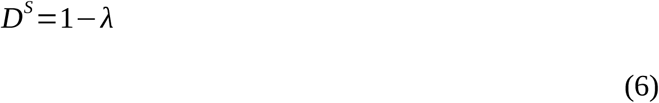

where *n*_*i*_ is the frequency of each *i* brewing yeast strain in a given beer category and *N* is the number of recipes for a beer category. Finally, the evenness of a specific brewing yeast strain among different beer categories was determined by the Pielou index (*J*) (Thukral, 2017) as follow (equation 7):

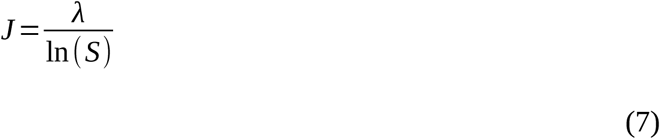

where λ is the diversity Simpson’s index as described in equation 5 and *S* indicates the total number of brewing strains for each beer category.

## 3. Results and Discussion

### 3.2. Beer categories and commercial brewing yeast strains data analysis

The craft beer revolution is a well characterized movement inside beer industry that can be roughly defined as the origin, development, and spread of local microbreweries as the consequence of the large-scale, homogeneous mildly beer brands that dominated the beer market in the late 20^th^ century followed by the increasing demand of new beer styles (Garavaglia & Swinnen, 2017). In addition, the craft beer industry can also be defined by consumers that drink less beer but are willing to pay more for special and pricier beers with different textures and flavors (Donadini & Porretta, 2017). Thus, it becomes clear that the major force that drives the craft beer revolution is the development of beer categories with a high diversity in the use of ingredients, where beers produced with local raw materials and yeast characterize the so called “beer du terroir” (Budroni et al., 2017). As pointed by Budroni et al. (2017), the use of local yeast strains or even the development of tailor-made yeast strains by different cellular and molecular techniques (Cubillos et al., 2019; Gibson et al., 2017) is a relatively unexplored tool for the diversification of local beers.

However, and despite the academic or industrial initiatives to promote the use of local ingredients, yeast manufacturers still have a major role in providing the main yeast strains used in breweries and then directly impacting the beer quality that is consumed. In order to understand the roles and the influence of commercial yeast strains in the fermentative aspects of different beer categories, the Brewer’s Friend, a large repository of beer recipes and yeast strain data was chosen to evaluate the specific parameters related to yeast strain richness and diversity as well as the preference of producers in the use of specific yeast strains for fermentation. It should be noted that beer categories that use lactic acid bacteria and non-conventional yeast genera and species (e.g., *Brettanomyces bruxellensis*) were excluded from this work. Thus, a total of 121,528 beer recipes divided into 34 major beer categories were downloaded from Brewer’s Friend web page. In addition, 476 commercially available yeast strains were analyzed in terms of type (ale or lager), formulation (dry or liquid), minimum and maximum fermentation temperature, and attenuation.

The data collected from Brewer’s Friend website showed that 14 beer categories have a high number of yeast strains in comparison to the average number of yeast strains employed for brewing (μ = 144.44 unique yeast strains × beer category^-1^, Figure 1A) and include relevant specialty craft beers, like India Pale Ale (IPA), Standard American Beer, Pale American Beer, and Belgian and Strong Belgian Ales (Figure 1A) (Haugland, 2014; Poelmans & Swinnen, 2018).

**Figure 1.**
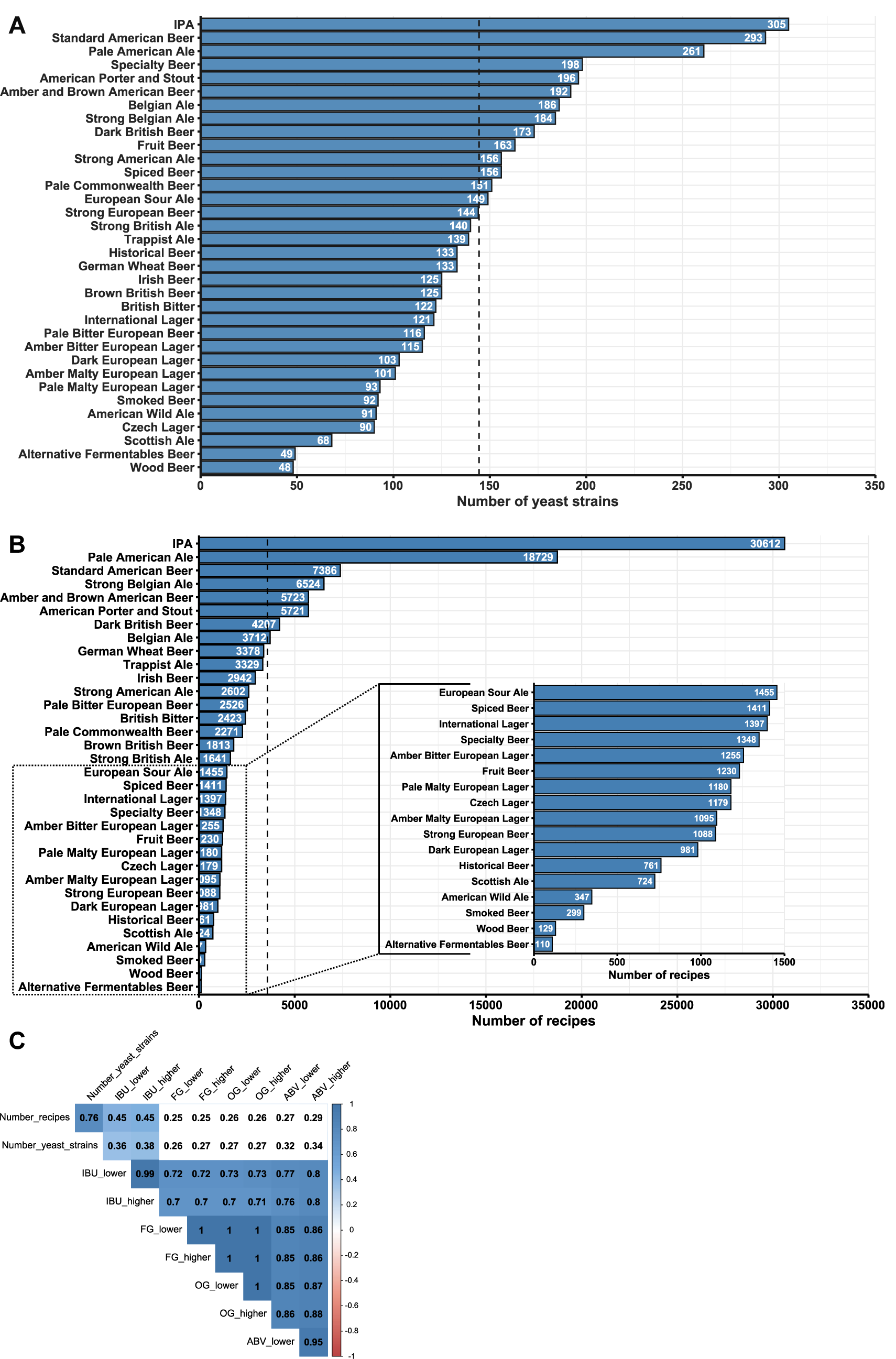
Determination of the number of unique yeast strains (A) and recipes (B) by each beer category. The dashed line in the graphics (A) and (B) indicates the mean value of the number of yeast strains and recipes, respectively. An amplified view of specific beer categories is indicated by the inset and dotted lines in graphic (B). In (C), correlation analysis of the number of yeast strains and recipes with major parameters associated to beer categories, like the lower and higher values of international bitter units (IBU_lower and IBU_higher, respectively), final gravity (FG_lower and FG_higher), original gravity (OG_lower and OG_higher), and alcohol by volume (ABV_lower and ABV_higher). The color scale in (C) indicates the pattern of correlation (negative or positive), from -1 (red) to 1 (blue).

Additionally, eight beer categories with a high number of yeast strains (IPA, Standard American Beer, Pale American Beer, Belgian and Strong Belgian Ales, American Porter and Stout, Dark British Beer, Amber and Brown American Beer) also display a high number of recipes compared to the average number of recipes by beer category (μ = 3574.35 beer recipes × beer category^-1^, Figure 1B). By its turn, the number of yeast strains and recipes used for beer categories related to lager family or specialty beers is low (Figures 1A and B). This initial data analysis prompted to question if the different beer categories parameters (e.g., IBU, OG, FG, and ABV) correlate with the number of yeast strains and recipes (Figure 1C). In fact, the number of yeast strains and recipes observed for a specific beer category did not show any correlation with OG, FG, and ABV level; however, it was observed a significant correlation of IBU level with the number of yeast strains and recipes (Figure 1C). This correlation could be partially explained by the increasing consumer preference for hoppier beer as well as the development of new hop cultivars that aggregate different flavors to the beer (Gabrielyan et al., 2014; Madsen et al., 2020), and thus directing the brewer’s preference for the design of beer recipes that made use of high amount of hops for bitterness or flavor. Moreover, a significant correlation of IBU, OG, FG, and ABV was also observed (Figure 1C).

Considering the total number of yeast strains evaluated in each beer category (Figure 1A), it was asked how many distinct yeast ale and lager strains in dry or liquid formulations were employed by the brewers in different beer categories (Figures 2A and B). From the total number of yeast strains annotated, it was observed that the number of liquid yeast strains counted for each beer category was higher than the number of dry yeast formulations (Figure 2A). Additionally, the number of distinct yeast ale strains determined for each beer category was higher than the number of lager strains (Figure 2B). These data could be supported by the fact that the number of commercially available yeast liquid formulations is higher than dry yeasts as observed from Brewer’s Friend website data (397 liquid versus 79 dry yeast strains) and the number of ale strains commercially available is also higher than lager strains (390 ale versus 86 lager yeast strains). An explanation about why there are many more liquid strains in comparison to dry strains (and the same for ale versus lager strains) was not completely addressed until now, but it can be hypothesized that many brewing yeast strains have a low tolerance to the industrial drying process, despite the fact that different methods to dehydrate yeast cells have been developed since the 18^th^ century (Gélinas, 2019). Supporting this hypothesis, it has been reported that lager yeasts strains have different desiccation tolerances (Layfield et al., 2011).

**Figure 2.**
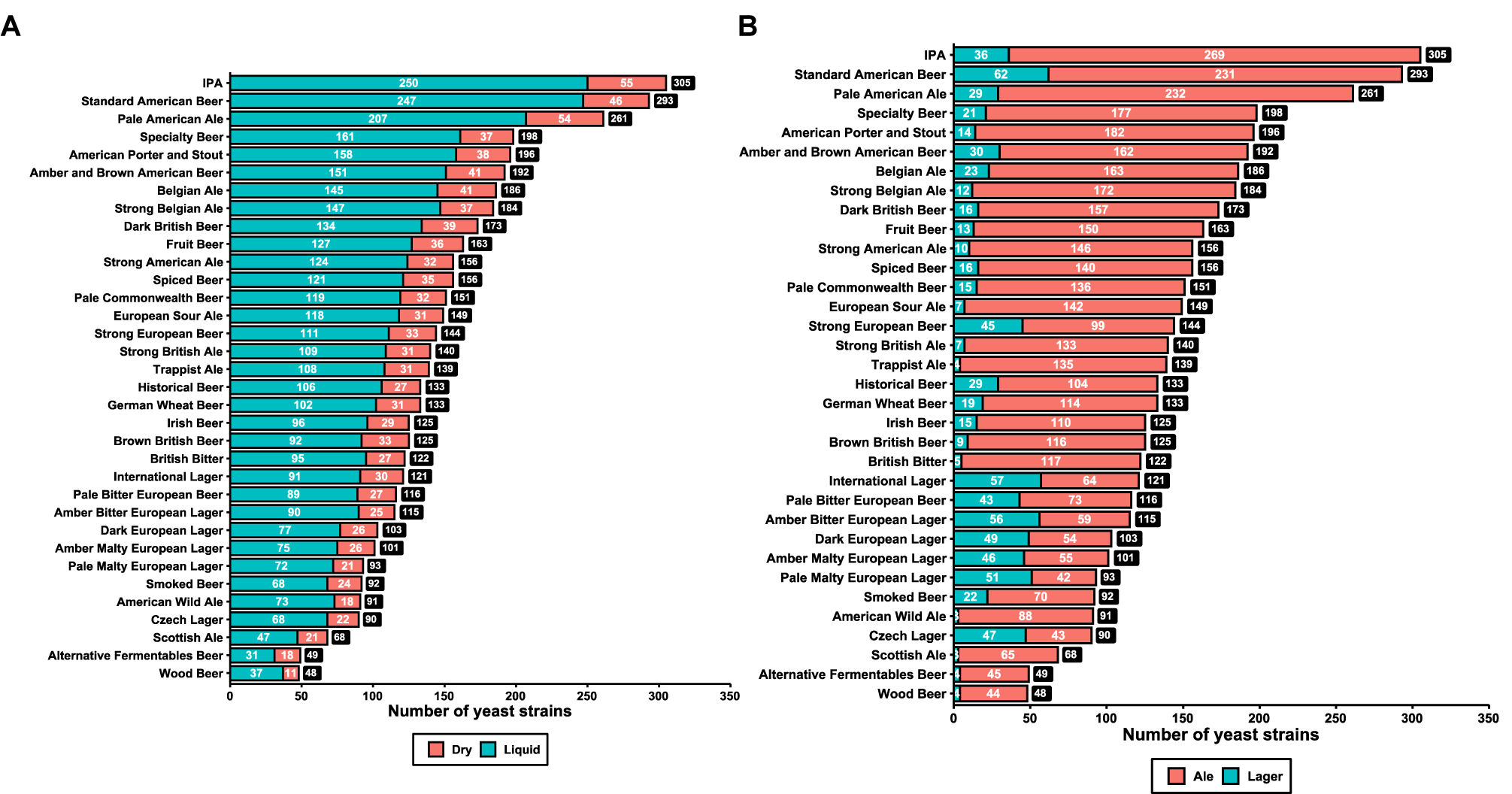
Number of dry or liquid (A) and ale or lager (B) yeast strains observed in each beer category. The total number of yeast strains in each beer category is indicated by the dark square in each column.

Thus, how similar are the commercial available brewers’ yeast strains in terms of fermentation temperature and attenuation? Considering the lower fermentation temperature reported by the yeast manufacturers for the 390 unique ale strains (68 dry and 322 liquid yeast formulations) used in brewing, it was observed that dry yeast strains have a significant lower mean fermentation temperature (μ = 16.42 °C; Figure 3A) in comparison to liquid yeast formulations (μ = 18.52 °C; Figure 3A). On the other hand, no significant difference was observed in the higher fermentation temperature reported by yeast manufacturers for ale dry (μ = 24.39 °C) and liquid (μ = 24.33 °C) formulations (Figure 3B). By its turn, from the 86 commercially available lager yeast strains (11 dry and 75 liquid formulations), it was not observed any significant difference in the mean lower fermentation temperature for lager dry (μ = 10.51 °C) and liquid (μ = 10.10 °C) strains (Figure 3C), while a significant difference was observed in the mean higher fermentation temperature for lager dry (μ = 17.85 °C) and liquid (μ = 14.52 °C) formulations (Figure 3D). Data regarding attenuation showed that ale yeast strains are similar in both dry (μ = 76.95%) and liquid (μ = 75.85%) forms (Figure 4A), while dry lager strains are significantly more attenuative (μ = 79.18%) than liquid strains (μ = 74.00%) (Figure 4B). To date, this is the first study that compared the attenuation and the higher and lower fermentation temperature of commercially available yeast ale and lager strains in dry and liquid formulations.

**Figure 3.**
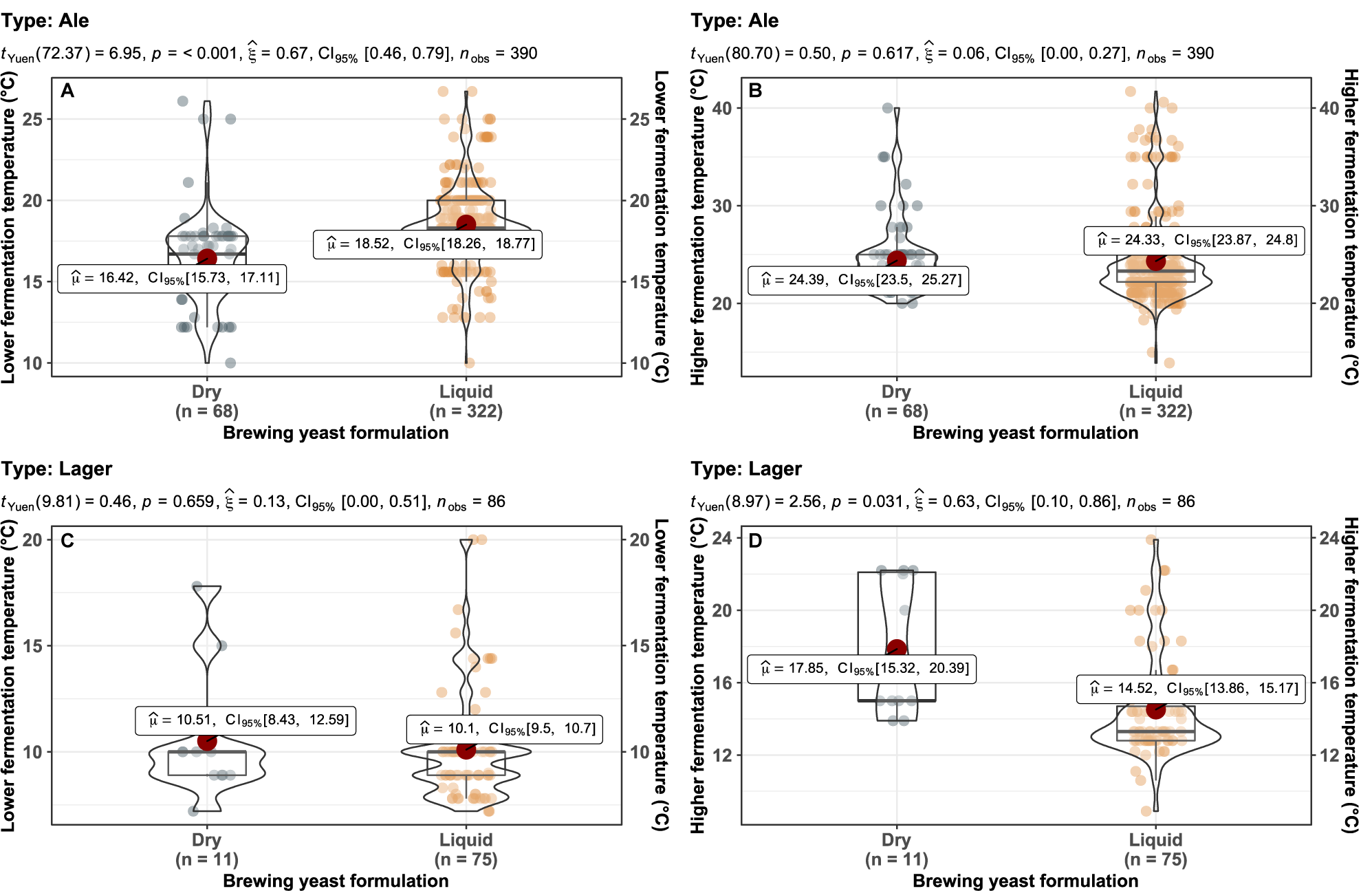
Evaluation of lower and higher fermentation temperatures (°C) for different ale (A, B) and lager (C, D) yeast strains (type) commercially available for brewers. The number of yeast strains observed for each formulation (n) as well as the statistical data analysis are indicated in the graphics.

**Figure 4.**
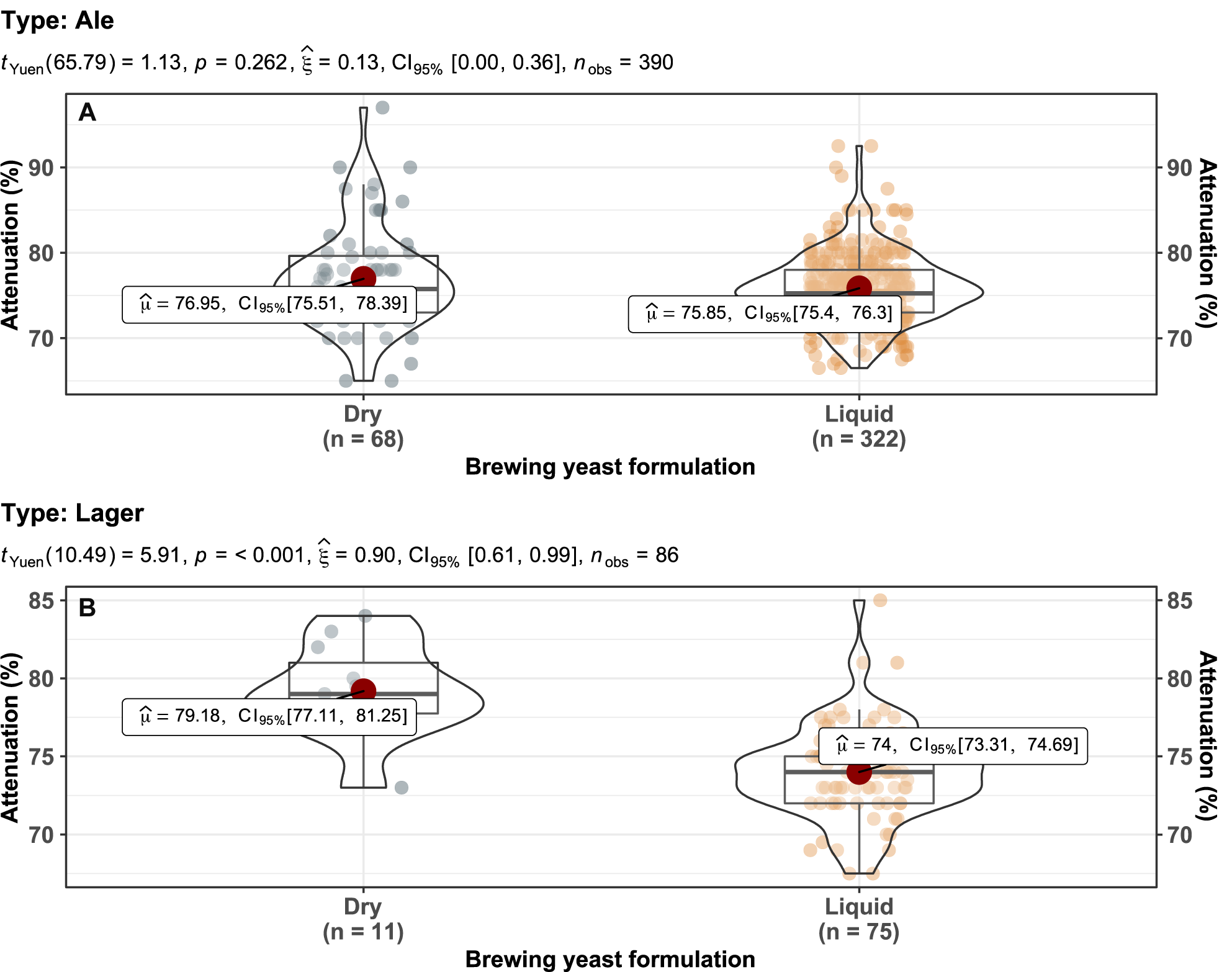
Evaluation of attenuation percentage for different ale (A) and lager (B) yeast strains (type) commercially available for brewers. The number of yeast strains observed for each formulation (n) as well as the statistical data analysis are indicated in the graphics.

Attenuation and fermentation temperature are the two main variables that significantly impact the beer, where the efficient use of malt-derived sugars by yeast strains (resulting in high ethanol yields) and the absence of off-flavors is desirable for any beer category (Powell et al., 2003). In this sense, it becomes clear from the data collected for this study that ale and lager yeasts strains in dry and liquid formulations are phenotypically similar considering the 95% confidence intervals (CI_95%_) for temperature (Figures 3A to D) and attenuation (Figures 4A and B). However, some outliers could be observed in ale strains with high fermentation temperature profiles (Figures 3A and B) which correspond to norwegian kveik and belgian hybrid saison strains (González et al., 2008; Preiss et al., 2018) as well as Kölsch/Altbier-associated yeast strains. By its turn, high fermentation temperature profile in lager yeast was observed for strains employed in the California Common beer style (Figures 3C to D). Regarding attenuation, the outliers found in ale strain data correspond to different belgian yeast strains that express the *STA1* gene (Krogerus & Gibson, 2020), leading to beer overattenuation (Figure 4A).

Fermentation temperature control is critical for many beer categories, as the yeast performance and the development of specific flavors are directly linked to fermentation temperature, especially modulating the production of esters and higher alcohol (Olaniran et al., 2011; Pires et al., 2014). Considering the data heterogeneity of yeast strains and recipes by each beer category analyzed (Figures 1A and B), what is the impact of brewer’s preference on lower and higher fermentation temperature for a given beer category? To answer this question, a weighted arithmetic mean value for the lower and higher fermentation temperatures (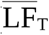 and 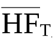, respectively) for each beer category was determined (Figure 5). Interestingly, two major groups of beer could be discriminated by considering the mean value of 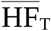 (μ = 22.56 °C), which were defined as “cold fermented beer” 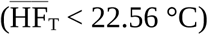 and “hot fermented beer” 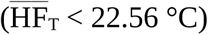 (Figure 5). The cold fermented beers correspond to all lager family-associated beer categories, while the hot fermented beer group contains all ale family-associated categories (Figure 5). A strong and positive correlation could be observed between 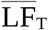 and 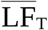, where the Czech Lager and Strong Belgian Ale categories correspond to the extremes of 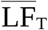 and 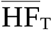 values (Figure 5).

**Figure 5.**
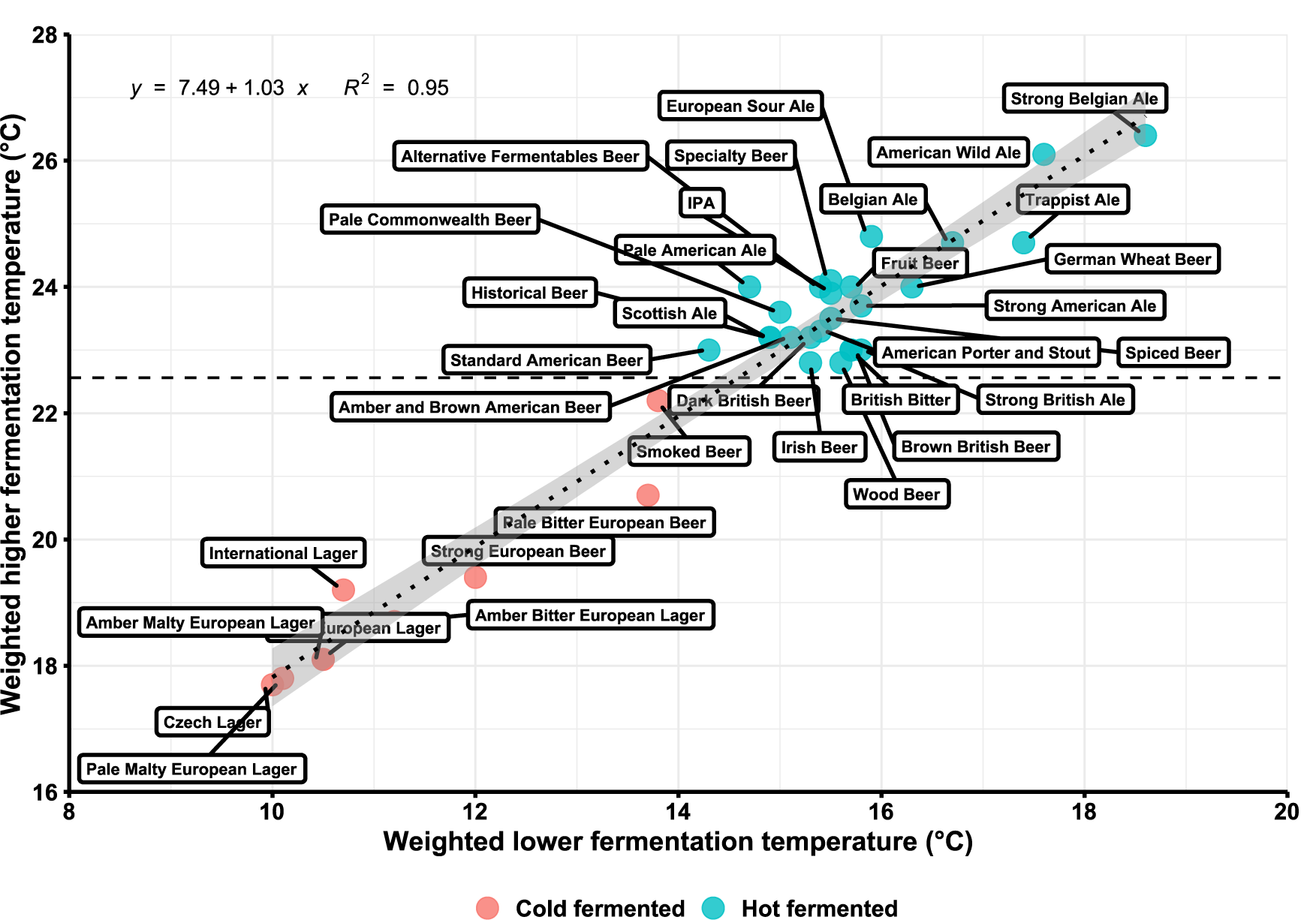
Linear regression of weighted lower and higher fermentation temperature (°C) observed for each beer category. The dashed line indicates the average value for weighted higher fermentation temperature. The coefficient of determination (*R*^*2*^) and the equation of linear regression are indicated in the figure. The dotted line and the respective the gray area indicates the regression line and confidence intervals, respectively.

### 3.2. The brewers’ preferences for yeast strain usage

The use of dry yeast is gathering popularity over yeast liquid formulations for brewing due to the fact that dry formulations occupy smaller volume and do not need refrigeration in comparison to liquid yeasts, resulting in lower costs associated with logistic and yeast storage. Moreover, dry yeast formulations can be kept for many years without loss of vitality (Rapoport, 2017). Thus, there is a natural tendency of brewers to employ dry yeasts in beer fermentation, despite the low number of ale and lager dry strains commercially available (68 and 11 strains, respectively). Interestingly, the preferential use of dry yeast rather than of liquid yeast for beer fermentation could be clearly observed from Brewer’s Friend data (Figure 6). The preferential yeast usage or *P*_*Y*_ was higher in all beer categories where dry ale and lager yeast strains were employed, while liquid ale and lager formulations were less preferred (Figure 6). Some hot fermented beer categories, like Standard American Beer, American Porter and Stout, Pale American Ale, Pale Commonwealth Beer, and IPA have high *P*_*Y*_ values for dry ale yeast strains (Figure 6). High *P*_*Y*_ values for dry lager strains were also observed for all cold fermented beer categories, with the exception of Pale Bitter European Beer category, which has a preferential use for liquid ale strains (Figure 6). Considering the brewer’s preferential yeast usage (Table 1), how this variable impacts the evenness, richness, and diversity of beer categories?

**Table 1.** Classification of major beer categories into fermentation types and analysis of evenness, richness, diversity, and preferential yeast usage.

| Category | Fermentation type | Evenness (Pielou index) | Richness (Menhinick index) | Diversity (Simpson index) | Preferential yeast usage (Py) |
| --- | --- | --- | --- | --- | --- |
| Smoked Beer | Cold fermented | High | High | High | Ale Dry, Lager Dry |
| Amber Bitter European Lager | Cold fermented | High | Low | High | Lager Dry |
| Amber Malty European Lager | Cold fermented | High | Low | High | Lager Dry |
| Strong European Beer | Cold fermented | Low | High | High | Ale Dry, Lager Dry |
| Dark European Lager | Cold fermented | High | Low | Low | Lager Dry, Lager Liquid |
| Czech Lager | Cold fermented | High | Low | Low | Lager Dry |
| Pale Malty European Lager | Cold fermented | High | Low | Low | Lager Dry |
| International Lager | Cold fermented | Low | Low | Low | Lager Dry |
| Pale Bitter European Beer | Cold fermented | Low | Low | Low | Ale Dry, Ale Liquid, Lager Dry |
| American Wild Ale | Hot fermented | High | High | High | Ale Dry |
| Historical Beer | Hot fermented | High | High | High | Ale Dry |
| Strong British Ale | Hot fermented | High | High | High | Ale Dry |
| Wood Beer | Hot fermented | High | High | High | Ale Dry |
| Brown British Beer | Hot fermented | High | Low | High | Ale Dry |
| Trappist Ale | Hot fermented | High | Low | High | Ale Liquid |
| European Sour Ale | Hot fermented | Low | High | High | Ale Dry |
| Fruit Beer | Hot fermented | Low | High | High | Ale Dry |
| Specialty Beer | Hot fermented | Low | High | High | Ale Dry |
| Spiced Beer | Hot fermented | Low | High | High | Ale Dry |
| Belgian Ale | Hot fermented | Low | Low | High | Ale Dry |
| Strong Belgian Ale | Hot fermented | Low | Low | High | Ale Dry |
| American Porter and Stout | Hot fermented | Low | Low | High | Ale Dry |
| Alternative Fermentables Beer | Hot fermented | High | High | Low | Ale Dry |
| Scottish Ale | Hot fermented | High | Low | Low | Ale Dry, Ale Liquid |
| British Bitter | Hot fermented | High | Low | Low | Ale Dry |
| Standard American Beer | Hot fermented | Low | High | Low | Ale Dry, Lager Dry |
| Amber and Brown American Beer | Hot fermented | Low | Low | Low | Ale Dry |
| Dark British Beer | Hot fermented | Low | Low | Low | Ale Dry |
| German Wheat Beer | Hot fermented | Low | Low | Low | Ale Dry, Ale Liquid |
| IPA | Hot fermented | Low | Low | Low | Ale Dry |
| Irish Beer | Hot fermented | Low | Low | Low | Ale Dry |
| Pale American Ale | Hot fermented | Low | Low | Low | Ale Dry |
| Pale Commonwealth Beer | Hot fermented | Low | Low | Low | Ale Dry |
| Strong American Ale | Hot fermented | Low | Low | Low | Ale Dry |

**Figure 6.**
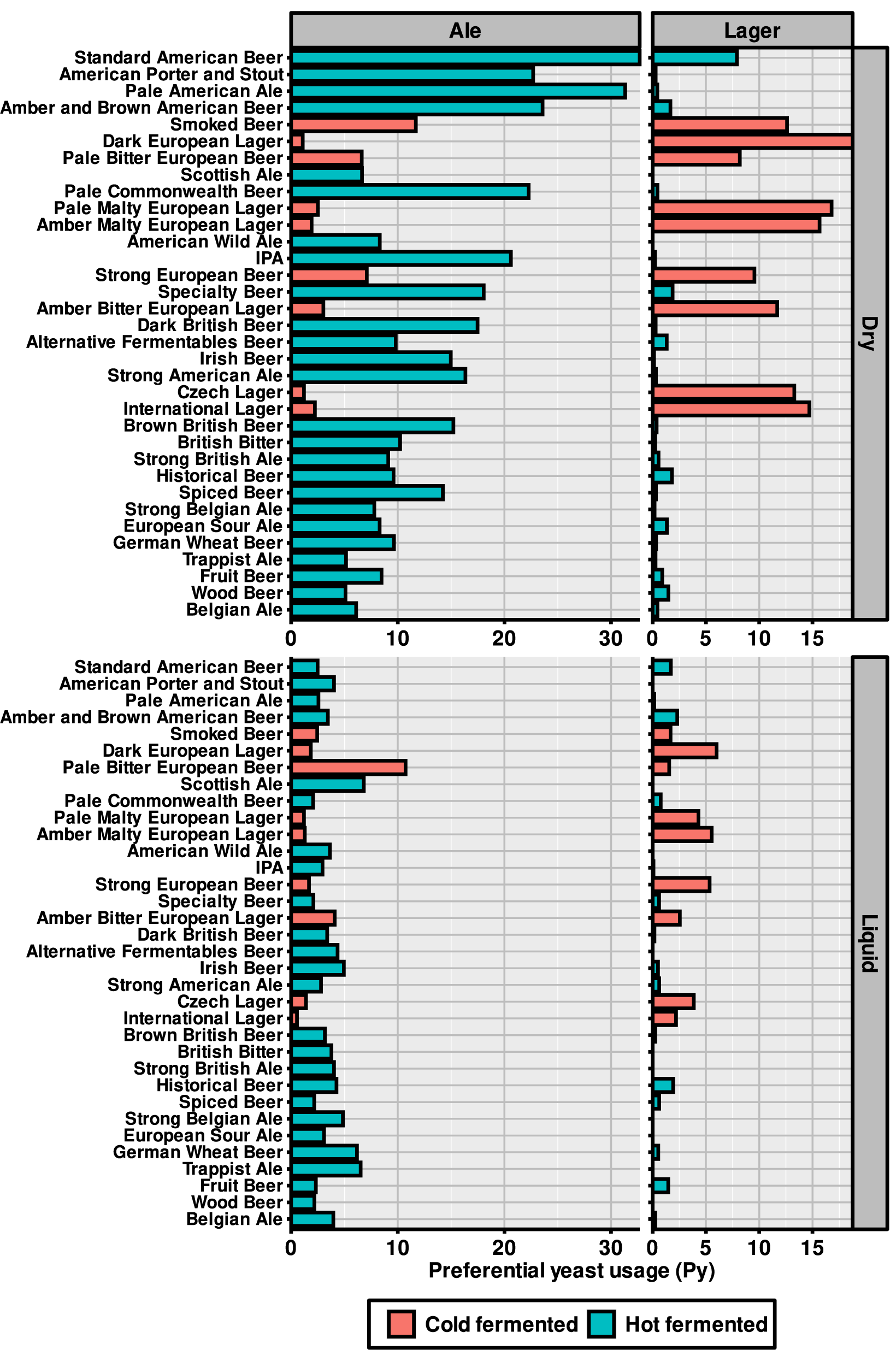
Preferential yeast usage (Py) analysis of ale and lager yeast strains in dry or liquid formulations for each beer category. Red bars indicate beer categories that are cold fermented, while blue bars indicate beer categories that are hot fermented.

### 3.3. Measuring the evenness, richness, and diversity distribution of commercial yeast strains in beer categories

To evaluate the impact of brewer’s preferential yeast usage in beer categories, a quantitative ecology analysis was performed. This analysis consider the concepts of evenness, richness, and diversity that are employed in different fields (Xu et al., 2020). For this work, the diversity concept is a variable that depends on the richness of different yeast strains found within a beer category, how evenness (homogeneous) are those strains distributed among beer recipes found in a category as well as the number of beer recipes found in a given category. Thus, a beer category with a high diversity has an elevated number of different yeast strains with an evenness distribution of those strains among a high number of beer recipes found within the beer category.

Initially, beer category diversity and richness were evaluated by using the Simpson’s (λ) and Menhinick (Mi) indexes, respectively, for cold and hot fermented beer (Figure 7A). By using the mean values of λ (μ_λCold_ = 0.93 and μ_λHot_ = 0.91) and *Mi* (μ_*Mi*Cold_ = 3.35 and μ_*Mi*Hot_ = 3.26) (Figure 7A) it was possible to classify beer categories into four major groups: (i) beer categories that have a high richness and diversity, (ii) beer categories with low richness and high diversity, (iii) beer categories with high richness and low diversity, and (iv) beer categories with low richness and low diversity (Figure 7A). A similar analysis was made considering λ diversity and Pielou evenness index (J),

**Figure 7.**
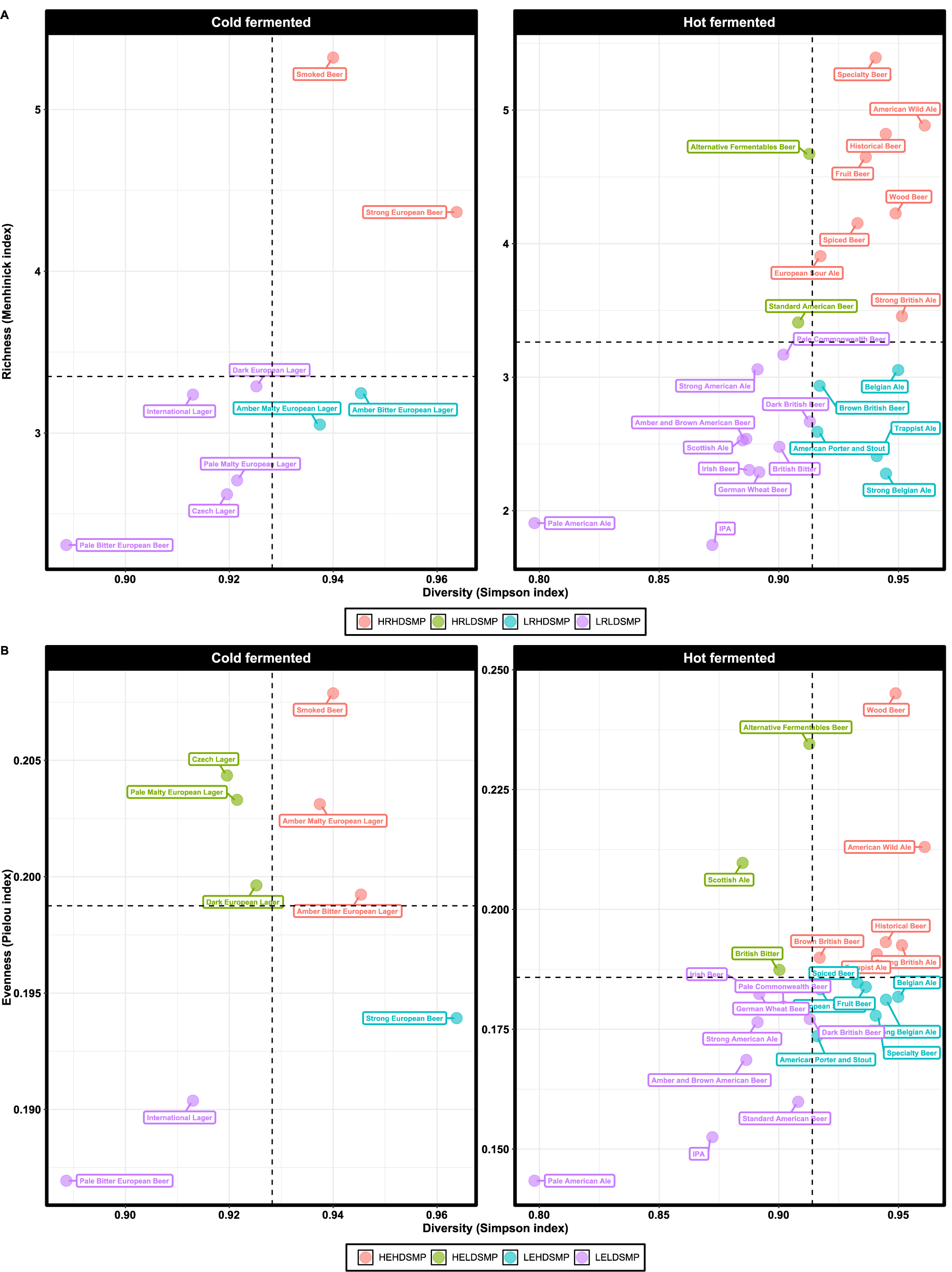
Analysis of brewing yeast strain richness and diversity (A), and evenness and diversity (B) for each beer category. Dashed lines indicate the average values of richness, evenness, and diversity. Abbreviations: High Richness-High Diversity_Simpson_ (HRHDSMP), High Richness-Low Diversity_Simpson_ (HRLDSMP), Low Richness-High Diversity_Simpson_ (LRHDSMP), Low Richness-Low Diversity_Simpson_ (LRLDSMP), High Evenness-High Diversity_Simpson_ (HEHDSMP), High Evenness-Low Diversity_Simpson_ (HELDSMP), Low Evenness-High Diversity_Simpson_ (LEHDSMP), Low Evenness-Low Diversity_Simpson_ (LELDSMP).

where the mean values for λ (μ_λCold_ = 0.93 and μ_λHot_ = 0.91; Figure 7A) and *J* (μ_*J*Cold_ = 0.19 and μ_*J*Hot_ = 0.18; Figure 7B) allow to group beer categories into four types: (i) high richness and evenness, (ii) low richness and high evenness, (iii) high richness and low evenness, and (iv) low richness and low evenness (Figure 7B). Considering cold fermented beer group, it was observed that Pale Bitter European Beer, Czech Lager, Pale Malty European Lager, International Lager, and Dark European Lager have low yeast strain diversity and richness (Figure 7A and Table 1), meaning that brewers preferentially use a small number of yeast strains, especially dry lager yeasts, to ferment beers that fall within these categories (Figure 6). Moreover, the evenness of yeast strains usage for Pale Bitter European Beer and International Lager is low (Figure 7B and Table 1), also pointing to a preferential use of yeasts type and formulation as seen in the previous analysis (Figure 6). On the other hand, Amber Malty European Lager and Amber Bitter European Lager have a high diversity and evenness, but a low richness (Figures 7A and B; Table 1), which can be explained by the extensive use of dry lager strains (Figure 6).

Noteworthy, from 25 hot fermented beer categories analyzed, ten categories display low values of richness, evenness, and diversity, like Pale American Ale, IPA, Strong American Ale, Amber and Brown American Ale, among others (Figures 7A and B; Table 1). This result indicates that brewers preferentially use a very low number of yeast strains to ferment beers that fall within these categories, corroborating the *P*_*Y*_ data that favor the use of dry ale formulations for these categories (Figure 6). Interestingly, Belgian Ale, Strong Belgian Ale, Trappist Ale, and Brown British Beer have a low richness and high diversity, pointing to the fact that the number of specific strains used in these categories is not high despite being evenly distributed (Figures 7A and B; Table 1). Additionally, the number of recipes described for these categories is also high (Figure 1A), which contributes to the diversity values observed for Belgian Ale, Strong Belgian Ale, Trappist Ale, American Porter and Stout, and Brown British Beer categories.

Specialty beers like Fruit Beer, European Sour Ale, Spiced or Wood Beers, and Strong British Ale have a high diversity and richness (Figure 7A and Table 1), indicating that brewers are prone to use a high diverse set of yeast strains to ferment beer that fall within these categories.

However, the evenness of yeast strain usage among these beer categories can be variable, where Fruit Beer, European Sour Ale, Strong British Ale, and Spiced beers display low evenness, while Wood Beer has a high evenness value (Figure 7B; Table 1).

## 4. Conclusion

The data gathered in this work showed that brewers have a preference for a small set of yeast strains, indicating that there is an underexplored potential for developing new beers by using the commercial yeast strains that are already available and have low usage. Despite the efforts of researchers to develop new yeast strains (Cubillos et al., 2019; Hittinger et al., 2018; Mertens et al., 2015; Saerens et al., 2010; Steensels et al., 2014), there is an ingrained brewing culture for using conventional yeast strains to ferment beer, especially dry ale formulations. As pointed before, dry yeast formulations have a series of advantages when compared to liquid yeast strains (Bellissimi & Ingledew, 2005), but the low number of dry yeast strains is a major disadvantage that brewers should consider on the development of new products. On the other hand, the low number of available dry yeast strains also indicates a potential and unexplored industrial field for the development of new dry yeast strains. For example, the design of strains with high biotransforming activity of hop-derived compounds is a major trend in brewing (Praet et al., 2012; Steyer et al., 2017; Tran et al., 2015) and can aggregate value to beer (Gabrielyan et al., 2014). Additionally, yeasts with increased resistance to osmotic pressure and high attenuation are gathering attention from brewers to develop new beers (Krogerus & Gibson, 2020).

In conclusion, yeasts are an underexplored resource in the brewing industry, with a large space for designing and repurposing commercially available yeast strains for the creation of new beers.

## Human and animal rights

The experiments included in this manuscript did not involve any animal or human participants.

